# Consecutive signaling pathways are activated in progression of Duchenne muscular dystrophy in *C. elegans*

**DOI:** 10.1101/532465

**Authors:** Heather C Hrach, Hannah S Steber, Jason Newbern, Alan Rawls, Marco Mangone

**Affiliations:** Molecular and Cellular Biology Graduate Program, School of Life Sciences 427 East Tyler Mall Tempe, AZ 85287 4501.; Virginia G. Piper Center for Personalized Diagnostics, The Biodesign Institute at Arizona State University, 1001 S McAllister Ave, Tempe, AZ, USA; Barrett Honors College, Arizona State University, 751 E Lemon Mall, Tempe, AZ 85281; School of Life Sciences 427 East Tyler Mall Tempe, AZ 85287 4501

## Abstract

**Background:** Duchenne muscular dystrophy (DMD) is a lethal, X-linked disease characterized by progressive muscle degeneration. The condition is driven by nonsense and missense mutations in the dystrophin gene, but the resulting changes in muscle-specific gene expression that take place in dystrophin’s absence remain uncharacterized, as they are potentially obscured by the chronic inflammation elicited by muscle damage in humans. *C. elegans* possess a mild inflammatory response that allows for the characterization of the transcriptome rearrangements affecting disease progression independently of inflammation.

**Results:** In effort to better understand these dynamics we have isolated and sequenced body muscle-specific transcriptomes from *C. elegans* lacking functional dystrophin at distinct stages of disease progression. We have identified two consecutively altered gene networks, which are also disrupted in the dystrophin deficient *mdx* mouse model. We found an upregulation of genes involved in mitochondrial function early in disease progression, and an upregulation of genes related to muscle fibre repair in later stages. This suggests that dystrophin may have a signaling role early in development, and its absence may activate compensatory mechanisms that counteract muscle degradation caused by loss of dystrophin. We have also developed a temperature-based screening method for synthetic paralysis that can be used to rapidly identify genetic partners of dystrophin.

**Conclusions:** Our results allow for the comprehensive identification of transcriptome rearrangements that potentially serve as independent drivers of disease progression and may in turn allow for the identification of new therapeutic targets for the treatment of DMD.

**One Sentence Summary:** A tissue specific transcriptome analysis of dystrophin deficient muscle in *C. elegans* reveals that dystrophin has distinct, dynamic signaling roles in early and late stage progression of Duchenne muscular dystrophy.

## BACKGROUND

Duchenne muscular dystrophy (DMD) is an X-linked, recessive disease caused by out of frame mutations in the dystrophin gene [1]. The dystrophin gene codes for a structural protein found beneath the sarcolemma, where it is anchored both to the dystrophin glycoprotein complex (DGC) and cytoskeletal actin, thus stabilizing the protein complex and the integrity of the cell membrane [2]. In humans, the absence of functional dystrophin results in progressive degeneration of the skeletal and cardiac muscles. The hallmark symptoms of DMD extend beyond muscle degeneration to include respiratory failure, cardiomyopathy, and pseudohypertrophy. The condition remains the most commonly diagnosed type of muscular dystrophy, affecting approximately 1 in 3,500 male births globally.

While the role of dystrophin in forming a physical connection between the extracellular matrix (ECM) and cytoskeleton has been well characterized [3], a comprehensive molecular definition of dystrophin’s function is not fully understood. Vertebrate models of DMD include the *mdx* mouse [4] and the golden retriever muscular dystrophy canine [5]. Both models have contributed significantly towards our understanding of dystrophin function, but they face limitations when studying the cell autonomous effects of dystrophin deficiencies on skeletal muscle independently of the myolysis and fibrosis associated with chronic inflammation observed in mammals. In addition, *mdx* mice do not breed well, making them difficult to use to study molecular mechanisms using approaches like genome wide screens and other large-scale studies [6, 7].

Mitochondrial dysfunction and systemic deregulation of cellular energy homeostasis have been observed in both in DMD patients and *mdx* mice [8-10]. Mitochondria are important players in this disease, as a steady-state flux of ATP, Ca+ and other components of energy metabolism are needed for muscle contraction. A recent study identified a significant increase in enzymes that are major consumers of NAD+ in dystrophic muscle tissues, and its replenishment was shown to ameliorate paralysis in *mdx* mice and *C. elegans* [8]. Evidence shows that mitochondria play a pivotal role in disease progression. However, it is still not clear if mitochondrial dysfunction is induced by the loss of muscle fibres, which in turn induce muscle paralysis and necrosis, or if it occurs in early stages of the disease when the paralysis and loss of muscle tissue has not yet been initiated [11].

The invertebrate model *C. elegans* has the potential to address these questions and serve as an informative model system for DMD. These nematodes possess a singular ortholog of the dystrophin gene (*dys-1*), which is similar in protein size to the human dystrophin gene, and contains similar actin binding and scaffolding regions, which are used to bridge the dystrophin complex to cytoskeletal actin (**Fig. 1A**). Additionally, several essential members of the DGC are conserved between humans and nematodes. The *C. elegans* DGC is largely comparable to the human complex, and includes key proteins such as dystrobrevin (*dyb-1*), sarcoglycans (*sgca-1, sgcb-1* and *sgn-1*), and the syntrophins (*stn-1* and *stn-2*). This predicts strong selective pressure for functional conservation of the DGC from *C. elegans* to humans.

**Fig. 1:**
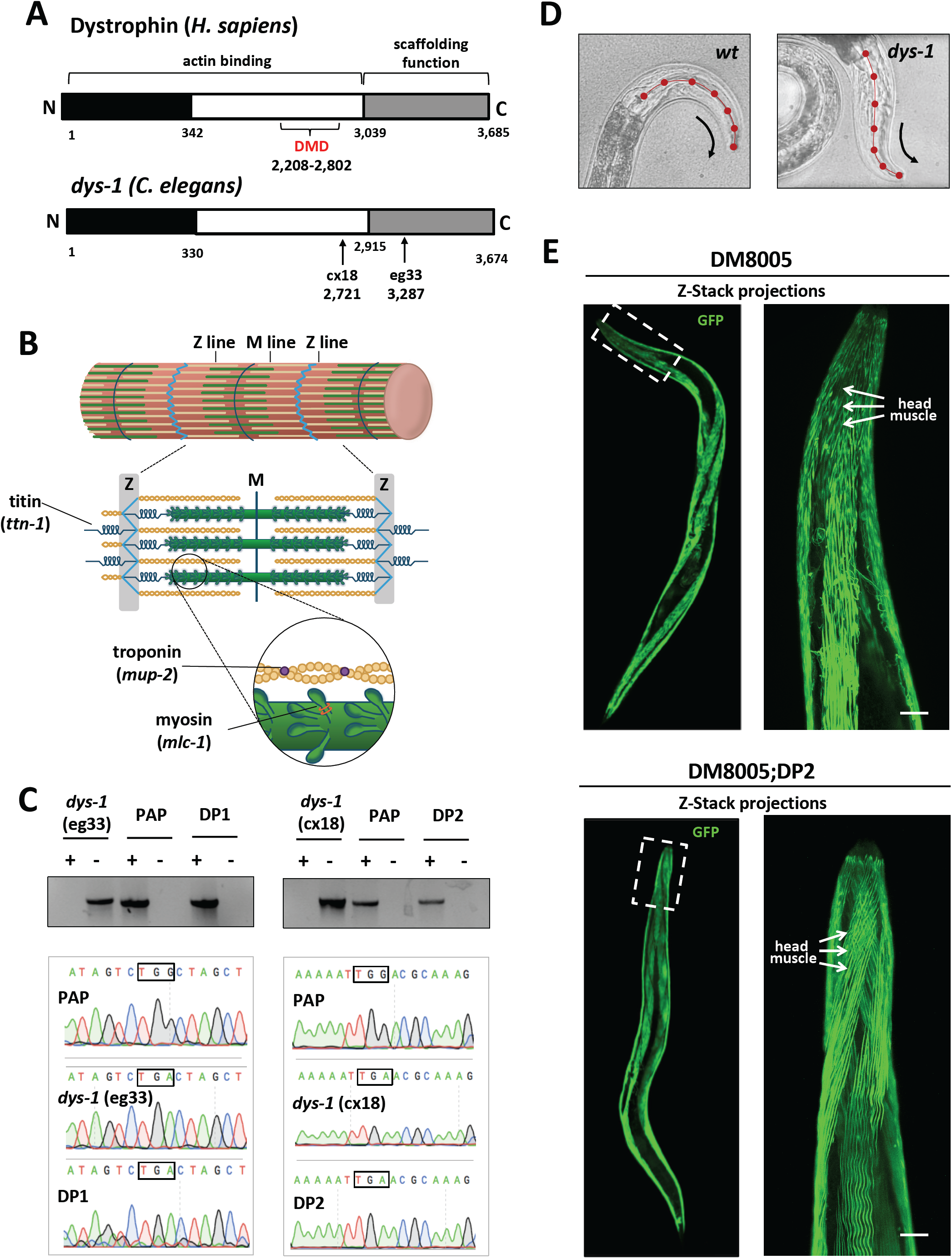
*C. elegans* as a model system for human DMD. (A) Diagram of functional protein domains in human dystrophin and the *C. elegans* ortholog *dys-1*. The functional domains of the human dystrophin protein are conserved in *C. elegans.* Mutational hotspot for DMD mutations in humans is labeled in red. The mutant strains *dys-1*(cx18) and *dys-*1(eg33) have nonsense mutations in the DMD ortholog *dys-1* in the actin binding and C-terminal scaffolding domains respectively. (B) A diagram of a human sarcomere highlights the human genes and associated *C. elegans* ortholog of several of the muscle structure related genes that were selected for RNAi screens. (C) DP1 and DP2 strains retain cx18 and eg33 mutations after being crossed with our PAP strain. Top panel: PCR analysis from genomic DNA extracted from *dys-1*(eg33), *dys-1*(cx18), DP1, DP2 and PAP strains. “+” denotes primers pairs that confirm the presence of a single copy integrated construct, and “-” denotes primers pairs that confirm the absence of an integrated construct. Lower panel: Trace files produced from the sequencing of genomic PCR of the *dys-1* locus confirm the presence of the nonsense mutation in our DP1 and DP2 strains. After crosses were performed, DP1 and DP2 strains retain both the associated nonsense mutations in the dystrophin gene, and the presence of our integrated PolyA Pull construct. (D) *dys-1* strains *dys-1(cx18) and dys-1(eg33)* exhibit a head bending phenotype. *wt* head bending coincides with direction of movement (black arrows), while *dys-1* head bending opposes direction of movement. (E) Confocal images of *wt* (DM8005, [myo-3p::GFP::myo-3 + rol-6(su1006)]) and dystrophin deficient strains ([myo-3p::GFP::myo-3 + rol-6(su1006)];DP1) for myosin heavy chain fibres. Right panels show an enlargement of the head muscles in both strains. There are no major changes in muscle structure between *wt* and dystrophic muscle in *C. elegans*.

*C. elegans* myogenesis has been extensively characterized [12], with only two main types of muscles: obliquely striated and nonstriated [13]. Many structural muscle proteins are also conserved between worms and humans, suggesting a maintained functional mechanism of contraction (**Fig. 1B**).

The *C. elegans* mutant strains *dys-1(cx18) and dys-1(eg33)* were recently introduced and represent a novel tool to study DMD *in vivo* [14, 15]. These mutant strains contain different nonsense mutations leading to the expression of a truncated DYS-1, which lacks the essential scaffolding portion of the C-terminus (**Fig. 1C**). *dys-1* gene expression is restricted to the worm body and vulva muscles. Mimicking the human condition, early reports have shown that loss of *dys-1* does not result in dramatic muscular degeneration phenotypes in juvenile worms. Instead they display defects in motility that are phenotypically distinct and associated with the loss of *dys-1.* This includes anomalous bending of the head, hyperactivity, and slight hyper contraction of the body wall muscle in early stages accompanied by an age-dependent, progressive loss of locomotor function (**Fig. 1D)**. In later stages, *dys-1* worm strains exhibit increased muscle cell death, attenuated motility and have shorter lifespans than *wt* worms (**Additional File 1: Figure S1**) [15].

Both *dys-1* worm strains display similar disease phenotypes, suggesting that both mutations, although in different portions of the *dys-1* gene, affect similar pathways. The introduction of human *dystrophin* cDNA in these mutant worms rescues these phenotypes [15], suggesting they are indeed a highly appropriate disease model.

*C. elegans* possess an innate immune response; the adaptive immunity is primitive and cell-mediated immunity is absent [16], leading to minimal muscle inflammation in *dys-1(cx18) and dys-1(eg33)* strains. The absence of chronic inflammation, which normally leads to myofiber necrosis and fibrosis in DMD patients or mouse models, causes these worm strains to move somewhat similarly to *wt* worms. Paralysis is subtle, localized to specific areas, and becomes most apparent in adult worms (**Fig. 1E**). Because the muscle tissues in these strains are not being actively damaged by chronic inflammation as in patients with DMD and *mdx* mice, this model is unique in its ability to study the cell autonomous contribution of changes in gene expression in the absence of muscle inflammation (**Fig. 1E**).

Although they were initially very promising, these two strains were not fully adopted by the community along with the *mdx* mouse, as a comprehensive analysis on the extent to which nematodes mimic the human version of the disease remained to be characterized. Additionally, these strains lacked a quantifiable phenotype that was easy to score, which initially prevented the development of large-scale genetic screens for the identification of therapeutic targets.

Our lab has previously adapted an established technique, which optimizes tissue-specific RNA extraction from intact organisms [17], allowing the identification of tissue-specific transcriptomes at single-base resolution [18-20]. This method, named PAT-Seq, takes advantage of the binding affinity and specificity of the cytoplasmic <u>P</u>oly<u>A B</u>inding <u>P</u>rotein (PABPC1) to polyA tails of mRNAs [21], which after UV crosslinking is immunoprecipitated, released from the tissue-specific polyA+ RNAs and the resultant transcripts are sequenced. *pab-1* is the worm ortholog of PABPC1, and by expressing this FLAG-tagged gene using MosSCI single copy integration technique in different worm tissues, our lab recently profiled *C. elegans* intestine, pharynx and body muscle tissues [18]. This study allowed us to define the body muscle transcriptome. We detected 2,610 genes expressed in this tissue, with only 329 unique genes corresponding to 365 spliced isoforms [18]. The list of top ten genes in the body muscle dataset was also enriched for muscle-specific genes such as myosin and actin isoforms. This study also detected a unique muscle-specific gene pool, including previously identified genes members of the *dystrophin* complex in human, such as dystroglycan, syntrophin, alpha-dystrobrevin, and dystrophin, the calveolin family, and type IV collagen.

In order to identify high quality cell-specific muscle specific transcriptome changes occurring in DMD, we crossed the *dys-1(cx18)* and *dys-1(eg33)* strains with our *myo-3p*::GFP::*pab-1*::3xFLAG worm strain (PAP) [18] (**Fig. 2A**), producing two novel strains named DP1 and DP2 respectively, which allowed the extraction and sequencing of high quality muscle-specific transcriptomes in worms lacking the dystrophin protein at different stages of the disease. This approach allowed us to fully define the dynamic transcriptome rearrangements occurring in the absence of inflammation at controlled time points, before and after the onset of paralysis. We have also performed targeted genetic experiments to study the contribution to paralysis of selected genes identified by our approach.

**Fig. 2:**
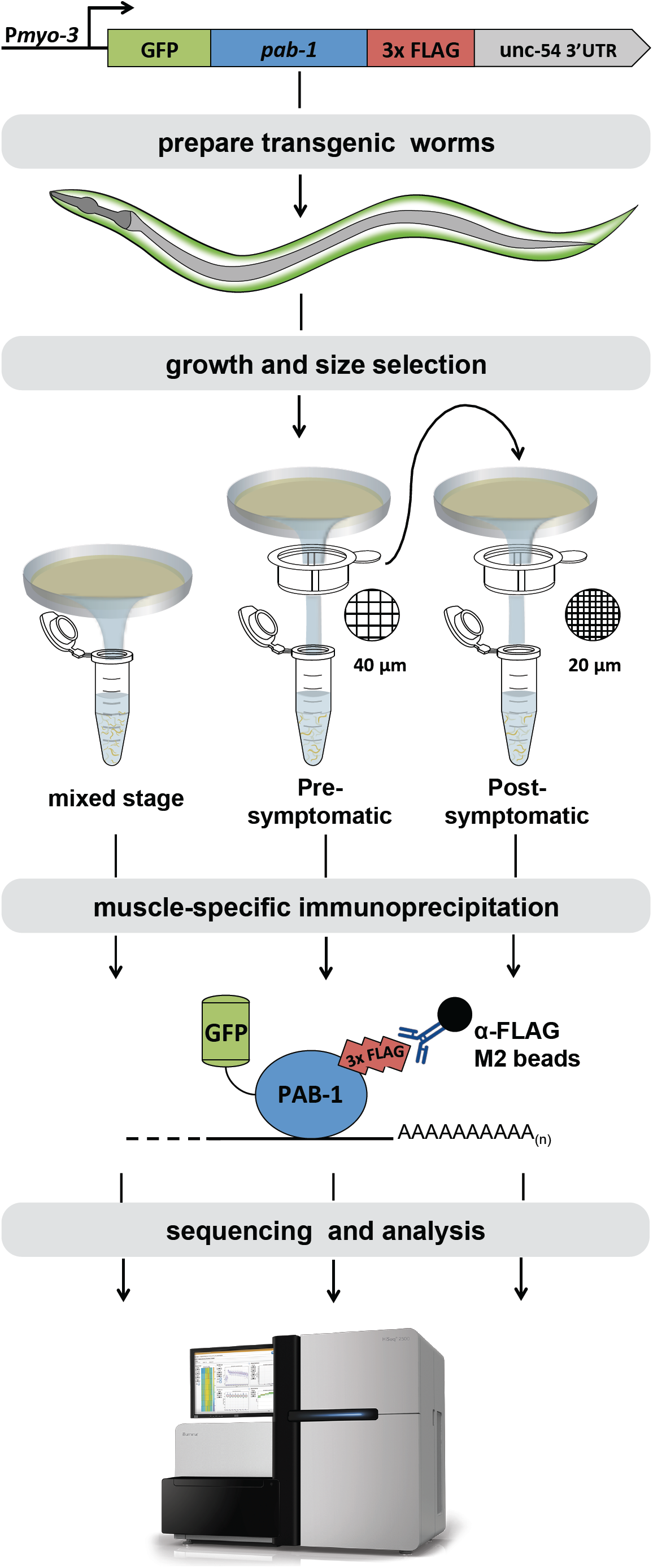
Pipeline used to isolate PRE and POST paralysis muscle specific transcriptomes. PAP expression construct places a FLAG-tagged *pab-1* under the control of a body muscle specific promotor *(Pmyo-3*) to isolate tissue-specific mRNA. PAP-expressing worms are crossed with *dys-1*(cx18) and *dys-1*(eg33) strains to establish new transgenic strains (DP1 and DP2) used in our muscle specific RNA-IPs. Worms were grown in solid culture on NGM plates and are sequenced directly for mixed stage IPs. The PRE and POST symptomatic datasets are obtained through sequential mechanical filtration. Muscle specific RNA-IPs are performed and sequenced in duplicate.

We have found that the absence of dystrophin first results in the deregulation of mitochondrial function prior to muscle paralysis. This impairment leads to differential expression of genes involved in muscle function and differentiation after the paralysis occurs perhaps as a part of a compensatory mechanism that is able to impede dystrophin-dependent muscle degeneration.

## RESULTS

### Pat-Seq from muscle tissues of *C. elegans* strains lacking a functional copy of the dystrophin gene produced high quality muscle specific RNAs

In order to perform PAT-Seq in *C. elegans* dystrophin deficient muscle tissues, we have taken advantage of the previously existing PAT-Seq technology used in our previous studies [18, 19]. The original PAP strain expresses the gene *pab-1* fused to GFP (N-terminus) and a 3xFLAG tag (C-terminus) restricted in the muscle tissue [18, 19]. *pab-1* is the *C. elegans* ortholog of the human cytoplasmic polyA binding protein (PABPC1), which typically binds the polyA track of mature mRNAs in the cytoplasm and is required for translation [22].

We have crossed the worm strain *myo-3p*::GFP::*pab-1*::3xFLAG (PAP) with *dys-1(cx18)* and *dys-1(eg33)* and prepared two new strains named DP1 and DP2. The two new strains retain the *dys-1* mutations cx18 and eg33, and express the muscle specific PAP transgene, which allows the muscle specific immunoprecipitation step using our *pab-1* based strategy. These two new strains (*dys-1(eg33)/pap* (DP1) and *dys-1(cx18)/pap* (DP2) were verified using Sanger sequencing to confirm the presence of the nonsense mutations in the *dys-1* gene in the crossed F1 worms that were GFP-positive in the muscle tissues (**Fig. 1C**). We also used a PCR approach to confirm the genomic integration of the PAP construct in the MosSCI locus [23] (**Fig. 1C**). In order to further validate our cross, we subjected these worm strains to a Kaplan-Meier survival analysis (**Additional File 1: Figure S1**). The average lifespan of N2 worms in this experiment is approximately 21 days, and both new strains behave very similarly to their reciprocal pre-crossed strain, with their lifespan drastically declining at day 18, with an overall increase in lethality when compared to *wt* worms (**Fig. S1**). Taken together these results suggest 1) we successfully crossed both dystrophin deficient strains with our PAP strain, and 2) the DP1 and DP2 strains reflect the phenotypes already characterized in the literature before and after our crosses, suggesting our PAP construct did not interfere with the *dys-1* phenotype.

We next performed the PAT-Seq experiments, isolating and sequencing muscle specific mRNAs from DP1 and DP2 strains. We wanted to detect precise changes in gene expression, not only in later stages of paralysis, but also before the paralysis was initiated, to have a more comprehensive overview of the dynamic gene changes occurring during the initiation and the progression of the disease. In addition, we wanted to reduce our sample number to simplify the data analysis and increase the depth of our sequencing results. Therefore, we decided to grow a large mixed population of our crossed worms, and then performed sequential mechanical filtrations by size, which led to the isolation of two pools of worms for each strain: one containing smaller and mildly phenotypic embryo-L2 worms, which we labeled ‘pre-symptomatic’ (PRE), and another containing larger and more extensively phenotypic L3-adult worms, which we labeled ‘post-symptomatic’ (POST) (**Fig. 2**).

We performed a total of 16 immunoprecipitations (see materials and methods), sequencing and analyzing DP1 and DP2 mixed stage samples, and PRE and POST paralysis samples in duplicate.

We obtained approximately 900M reads across all our sequenced samples, with approximately 50M reads each dataset (**Additional File 1: Table S1**). Within these samples we were able to map 42% of the reads to the *C. elegans* genome (WS250) (**Additional File 1: Table S1**). Both experimental and biological replicates in each dataset correlate well with each other (**Additional File 1: Figure S2**), with ∼2,000 shared protein coding genes within each group (**Additional File 1: Table S2**). The total and mapped reads for each sample are listed in **Additional File 1: Table S3**.

Our sequencing efforts detected a total of 3,748 genes across all datasets (**Additional File 1: Figure S3A**). Within this group 47% have been previously mapped in the body muscle [19] (**Additional File 1: Figure S3A**). When we repeated this analysis separately in our PRE and POST datasets, this percentage increases to ∼60% similarity (**Additional File 1: Figure S3A**), with >85% identity in our top 250 genes in all our datasets (**Additional File 1: Figure S3B**).

Taken together, these results suggest that our muscle specific RNA pull down was successful and able to identify *bona fide* muscle specific genes.

### Mitochondrial response is involved in the initiation of paralysis in our PRE dataset

The PRE symptomatic dataset is mainly composed of embryo to L2 mildly symptomatic worms. We obtained 1,932 protein coding genes expressed in this group, with 176 unique genes not expressed elsewhere (**Fig. 3** and **Additional File 1: Table S3**). There are very few genes that change drastically in expression level between the *wt* and PRE dataset (**Fig. 3**). A GO term analysis on the top 50 genes identified an enrichment in genes involved in nematode larval development and locomotion, such as *cah-4, dpy-30, ran-1, ceh-20, vab-15, ned-8, rpia-1, uri-1, tfg-1*, and *bus-8*.

**Fig. 3:**
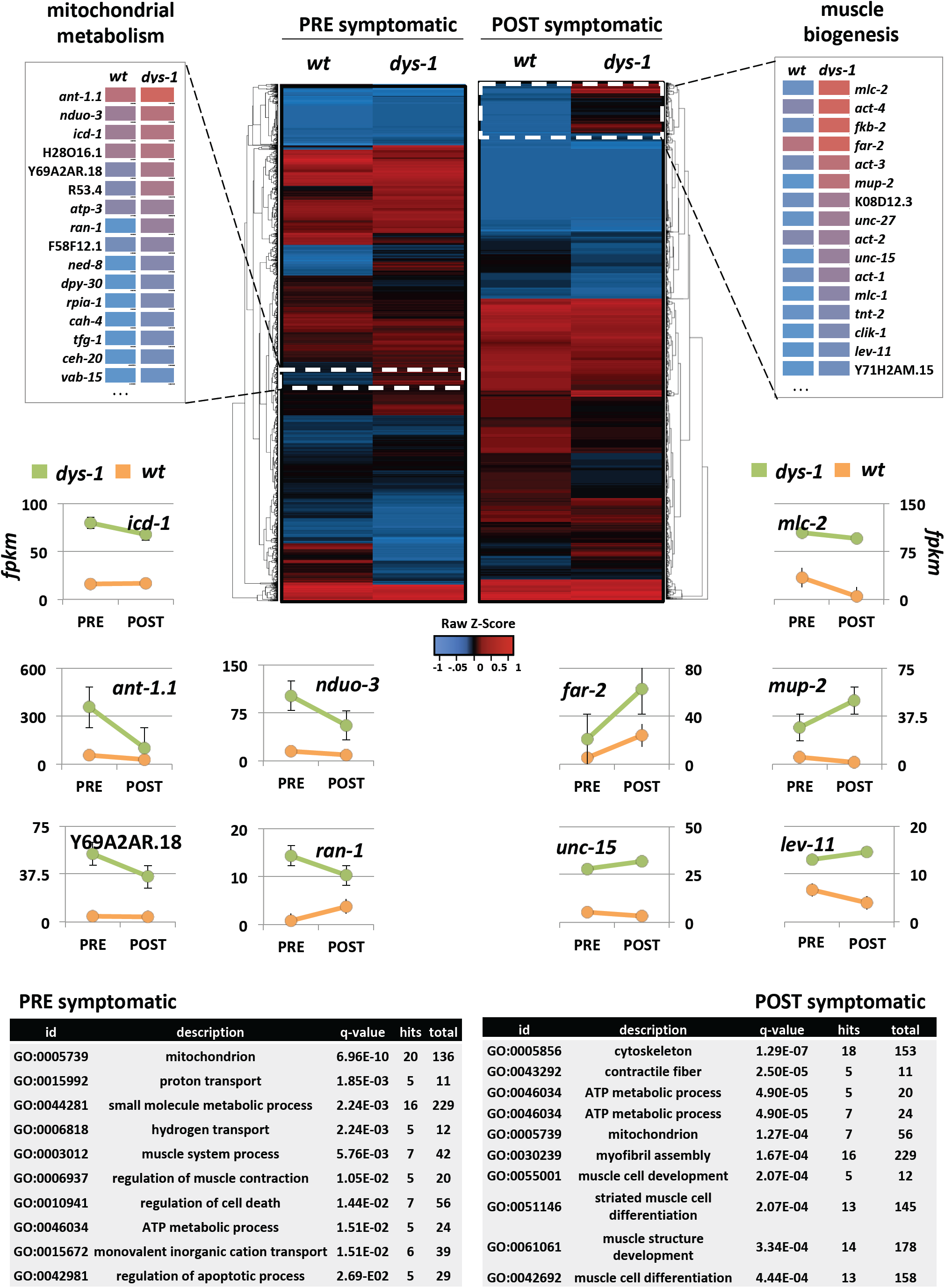
Summary of the muscle specific PAT-Seq results. The heat map summarizes the average changes in gene expression of both dystrophin deficient strains (DP1 and DP2) as compared to the *wt* strain. White boxes in the heat map mark genes that are upregulated in DP1 and DP2 strains in PRE and POST datasets. An expansion of each of these groups details relative change in expression level for genes selected based on function and rank in our datasets. Below heat maps the change in expression level of several genes are graphed linearly between PRE and POST symptomatic stages (green: median between DP1 and DP2; orange: wt). GO term analysis of the top 50 genes uniquely present either in our PRE or POST datasets shows an enrichment in genes involved in mitochondira metabolism (PRE) and muscle biogenesis (POST).

We also detect an unusual abundance of mitochondrial genes involved in ATP production and regulation of apoptotic processes within the top hits identified in our PRE dataset (**Fig. 3**). *ant-1.1* is an ADP/ATP translocase with a three-fold increase over the median between DP1 and DP2 datasets (when compared with the *wt* PRE dataset). This mitochondrial membrane receptor is responsible for transporting ATP synthesized from oxidative phosphorylation into the cytoplasm and absorbs back ADP in a 1:1 molar ratio [24, 25].

The genes Y69A2AR.18 (5-fold), F58F12.1 (3-fold), *icd-1* (2.5-fold), H28O16.1 (2-fold), *atp-3* (2-fold) are all mitochondrial ATP synthase subunits involved in the synthesis of ATP from ADP and inorganic phosphate.

Only 173 genes were uniquely detected in this stage that were absent in our *wt* control, and are most likely the direct result of the initiation of the disease. Most of these genes have an unknown function, but when aligned to the human proteome many of them show significant matches to known genes. *smo-1* is among those significantly abundant. *smo-1* is the *C. elegans* homolog of SUMO, a small ubiquitin-like signaling modifier that is attached to protein dictating localization and function.

### Genes involved in myogenesis and muscle contraction are overexpressed in the POST dataset

In our POST dataset we detected 2,273 protein-coding genes, which accounts for only 58% of genes in common with the PRE dataset (**Fig. 3** and **Additional File 1: Table S3**). This discrepancy is consistent with the progressive nature of the disease and the associated changes in muscle structure and overall health.

Essential constituents of the mitochondrial metabolism are also overexpressed at this stage, although at lower levels than in our PRE dataset. They include the mitochondria ATP synthase *Icd-1, ant-1.1*, H28O16.1, Y69A2AR.18, *atp-2* and *atp-3, nduo-1* (NADH-ubiquinone oxidoreductase), and W09C5.8 (cytochrome c oxidase).

Importantly, we detected an abundance of genes involved in myogenesis; including *mlc-1* and *mlc-2* (myosin light chain), *mup-2* (troponin), *act-1* (actin), *unc-27* (troponin), *lev-11* (tropomyosin), and *unc-15* (paramyosin). This result is surprising, as an increase in sarcomeric gene transcription is consistent with muscle hypertrophy or active replacement of muscle, which does not occur in *C.* elegans as they lack a satellite cell equivalent. In this context, the increase in transcription may suggest a hidden compensatory signaling pathway activated in the absence of *dys-1* that precedes cell death.

Only 38 genes are uniquely present in our POST dataset, and the majority of these genes have unknown functions.

### A novel synthetic screen to identify genetically linked *dys-1* targets

Our approach identified specific genes differentially regulated in both dystrophin deficient samples across replicates. These genes were categorized by function, and among several trends, we identified an enrichment in our pre dataset of genes involved in mitochondrial metabolism, and the establishment and maintenance of muscle structure and function in our POST dataset.

In order to validate and expand the biological significance of these results, we decided to perform a targeted genetic screen testing genes identified by PAT-Seq for their ability to enhance muscle damage in dystrophin deficient strains when knocked down through RNAi. We reasoned that if there were genetic compensatory mechanisms in place to counteract paralysis, the knock down of upregulated genes detected in our screen would lead to detectable impairment of muscle viability that would suggest a genetic link between *dys-1* and the tested genes.

The *dys-1* phenotype is subtle when compared to *wt* worms. For this reason, we decided to use a previously established strain that uses a background mutation to enhance paralysis to create a more definitive and scorable phenotype.

It has been previously shown that combining dystrophin mutations with mutations in the transcription factor MyoD exacerbates dystrophic phenotypes in mice, and that this combination has a similar effect in *dys-1* strains [26]. The temperature sensitive strain *hlh-1(cc561ts)* contains a hypomorphic mutation in the *C. elegans* homologue of MyoD, *hlh-1*, which renders these strains viable at the permissive temperature 15°C and severely uncoordinated with defects in body muscle morphology at non-permissive temperatures above 20°C [27].

We utilized this single mutant strain *hlh-1(cc561ts)*, as well as the double mutant strain *dys-1(cx18); hlh-1(cc561ts)*, and observed that at 15°C (permissive), both single and double mutant strains hatch and develop to adulthood without defects. In order to ameliorate the uncoordinated defects and lethality obtained at 20°C, we decided to decrease this temperature to 18°C (semi-permissive). At 18°C, ∼75% of the single mutants, and ∼50% of the double mutants are still viable (**Fig. 4A-B**), providing us with a distinct and tractable range to detect changes in incidence of paralysis following the RNAi experiments (**Fig. 4C**).

**Fig. 4:**
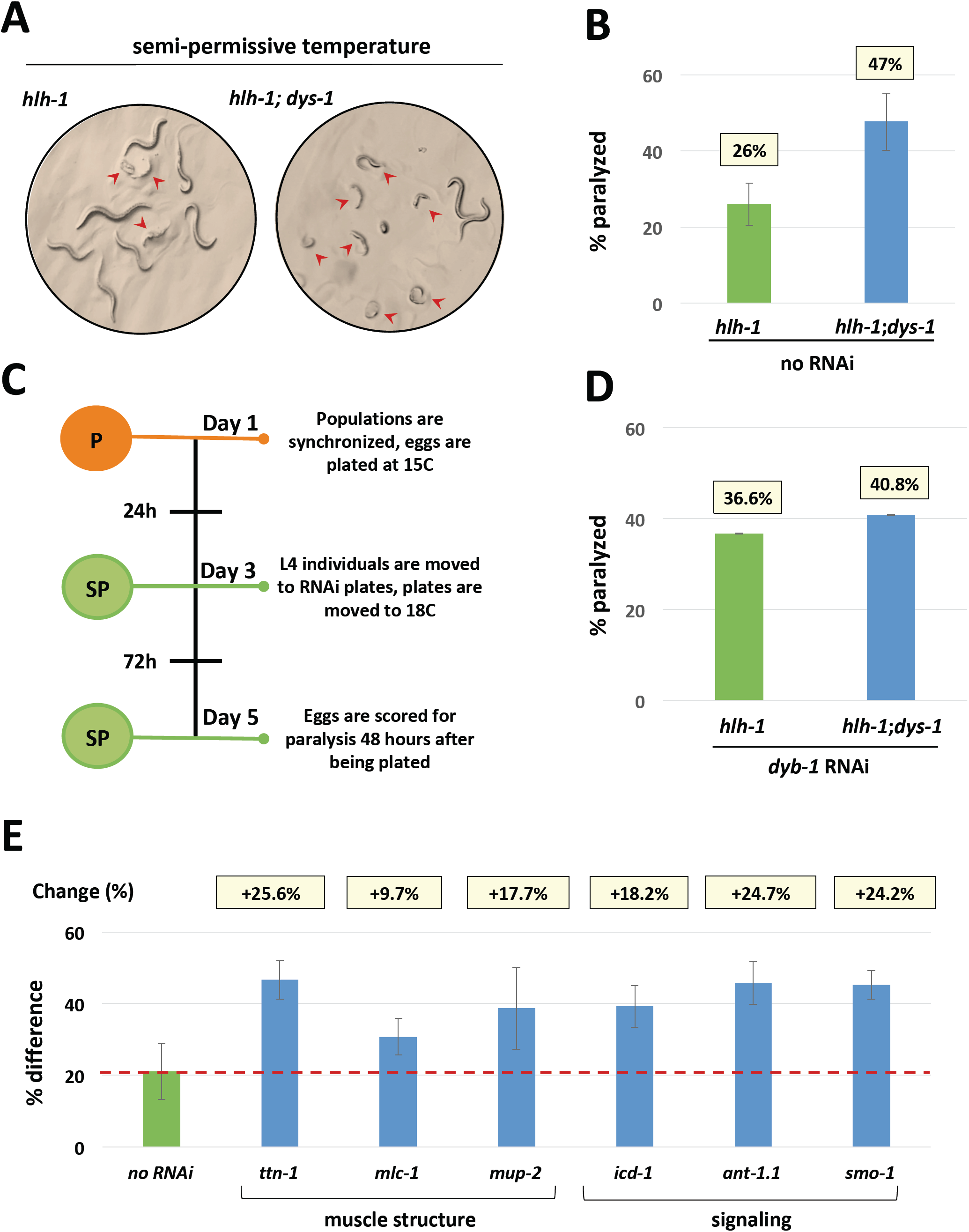
A synthetic screen to identify genetically linked *dys-1* targets. (A) Single *(hlh-1)* or double *(hlh-1; dys-1)* mutant sample populations scored for paralysis at semi-permissive temperature after 48 hours after parent progeny are plated. Red arrows indicate worms that were scored as paralyzed based on defects in morphology and resulting lack of movement (B) Quantification and comparison of average incidence of paralysis for single and double mutant strains at semi-permissive temperature. At this temperature *dys-1* is responsible for a 21% increase in paralysis in the presence of an *hlh-1* sensitized genetic background. Average % paralysis of 10 replicate plates summarized above each strain. A total of 543 worms were scored for semi-permissive experiments without RNAi. (C) Overview of semi permissive RNAi screen protocol describing which steps occur at permissive and semi-permissive temperatures. (D) Percentage of paralyzed worms following RNAi for *dyb-1,* with average % paralysis of 5 replicate plates summarized above each strain. (E) Synthetic paralysis RNAi assay. Each gene is displayed as the difference in paralysis observed between the *hlh-1* and *hlh-1;dys-1*(cx18) mutants and normalized to the difference of paralysis obtained from semi-permissive control screens (21%). Red dotted line indicates point of normalization for change in paralysis. Genes were grouped according to function (muscle structure and signaling). Percent increase change is shown on top of each bar chart.

Populations were synchronized and grown at 15°C (permissive) until they reached the L4 stage, and were then moved to a semi-permissive temperature of 18°C and allowed to lay eggs for 24 hours. At semi-permissive temperature, the single mutant strains *hlh-1* gave rise to progeny of which 26% developed with severe defects in body morphology and were almost completely paralyzed (**Fig. 4B**). Under the same conditions, the double mutant strains *hlh-1*; *dys-1* yielded a 21% increase in paralysis and severe defects in body morphology (**Fig. 4A-B**).

We then selected a subset of representative genes identified by PAT-Seq as being upregulated at different degrees, which are indicated with an asterisk in **Additional File 1: Table S3**. These genes were also chosen based on function, primarily because of their known roles in mitochondrial metabolism, muscle structure, and signaling function (**Fig. 4C**). We speculated that if these genes were indeed abundant in paralyzed worms as a compensatory mechanism to counteract paralysis, their depletion in the double mutant background would enhance paralysis when compared to the single mutant background.

The RNAi experiments were performed at the semi-permissive temperature of 18°C. Worms were scored for incidence of paralysis and morphological defects (**Fig. 4C**). The results between control and experimental strains were compared with each other, and then to the observed differences in semi-permissive experiments performed on single and double mutant strains in the absence of RNAi (**Fig. 4B**). As a control, we performed a knockdown of a known member of the nematode DGC, *dyb-1*. The knockdown of *dyb-1* was able to increase incidence of paralysis in the single mutant to an extent that reflected the double mutant in semi permissive control experiments (**Fig. 4D**).

We then tested our muscle and signaling pools. Although some of the genes were synthetic lethal in combination with the *hlh-1* background mutation (*act-1, cmd-1, unc-27* and *unc-15*), and could not been tested further, the majority of genes tested were able to enhance paralysis, although to different degrees (**Fig. 4E**) (**Additional File 1: Table S3**).

## DISCUSSION

Here we have described the use of the nematode *C. elegans* to study the cell autonomous molecular events associated with the functional loss of the dystrophin gene that leads to Duchenne muscular dystrophy in humans. We have performed genetic crosses to establish transgenic strains that serve both as a model for DMD, and as a functional tool to perform high quality, muscle specific RNA-IPs at a resolution that has not yet been achieved. Using these strains, we have sequenced and analyzed muscle specific transcriptomes at distinct developmental stages during disease progression, and identified several differentially regulated pathways in dystrophic nematode muscle.

We have also developed a novel, temperature-based genetic screen and performed a quantitative analysis to assess the functional significance of the identified genes in the viability of muscle cells, thus confirming their contribution towards DMD initiation and progression. This approach will allow a direct evaluation the role of specific genetic pathways in the clinical severity of DMD, which will be fruitful in developing drug targets for treating DMD patients.

Our dystrophin deficient strains DP1 and DP2 display phenotypes and altered molecular pathways that are very similar to those observed in *mdx* mice and DMD patients, strongly implying that this model system phenocopies many aspects of the disease at the molecular level.

These new strains recapitulate phenotypes previously characterized in the literature for *dys-1(cx18)* and *dys-1(eg33)*, with shortened average and maximum lifespan, hyperactivity and excessive bending of the head [14, 15] (**Additional File 1: Figure S1**), suggesting that introducing the PAP cassette into the *dys-1* genetic background has not altered the *dys-1* phenotype. In addition, the muscle morphology of both *dys-1* strains is highly similar to wild type muscle, as the muscle fibres do not present gross defects in morphology.

Our muscle specific sequencing approach in *C. elegans* allowed us not only to bypass the noise introduced by the immune response, as in *mdx* mice, but also to profile muscle tissue without contamination. Isolating pure muscle tissue from mice and patients is challenging, and although results are very informative and shed light on progression of DMD [28, 29], these samples are potentially contaminated with myofibroblasts, connective tissue, inflammatory cells, nerves and endothelial cells, and the results may be difficult to reproduce or validate due to intrinsic sample variation within experiments. Our approach bypassed these issues, providing the community with high-quality muscle-specific transcriptomes in the absence of functional dystrophin with minimal contamination and at single-base resolution.

Our study uncovered ∼ 2,000 protein coding genes present in early and late stage dystrophic muscle. Many of these genes are expressed in the muscle, and replicate what we already profiled in this tissue (**Additional File 1: Figures S3A-B**). Combined with a strong correlation observed between our biological replicates (**Additional File 1: Figure S3C**), these results suggest that our approach can accurately and consistently detect transcripts between biological replicates.

Because our approach is biochemistry-based, we were concerned about a high signal-to-noise ratio, which would have prevented the identification of fine genetic changes in our PRE and POST datasets. For this reason, we have applied a stringent bioinformatic filter to our datasets, restricting our analysis to the top 30-40% of the total genes identified in this study (**Additional File 1: Figure S3D**), which may have in turn lowered the number of genes considered, but provided us with higher quality results (**Additional File 1: Figure S3D**).

Our study revealed two distinct sets of genes that contribute to the DMD phenotype in *C. elegans*; the first set is activated before paralysis and participates in mitochondrial homeostasis, cell death and protein degradation signaling in the muscle, and the second set is activated at or after the onset of paralysis and is responsible for the establishment and maintenance of muscle structure.

Although there have already been studies in *mdx* mice that have reported the involvement of both pathways in DMD progression [9, 14, 30], it was still unclear how these two pathways are intertwined with each other, and in what order these molecular events occur. This question is important, as impaired mitochondrial metabolism has been already observed in *mdx* myoblasts in which a functional DGC has not yet been assembled [31], suggesting that these two pathways may be somewhat independent.

More precisely, it is still unclear if the damaged incurred by muscle tissue in absence of dystrophin leads to an increase of extracellular Ca2+ accumulation in muscle, which in turn causes mitochondrial membrane potential to collapse, or if the altered ATP synthesis in the mitochondria in the muscle tissue before paralysis causes muscle weakness, instability and degradation.

Our results suggest that these two pathways may not occur simultaneously; the mitochondrial dysfunction is detected early in disease progression, in accordance with previous findings [9], either before paralysis is initiated or while it is still mild.

In the PRE dataset we detected the differential regulation of many mitochondrial genes involved in glycolysis and ATP/ADP transport (**Fig. 3 and Additional File 1: Table S5**). Proteomics studies in *mdx* mice support these results [9]. Muscle contraction is dependent upon ATP production and usage, and its abundance is tightly regulated in healthy muscles.

Aberrant mitochondrial activity has been previously reported in patients and animal models with DMD [8, 32-36], even prior to dystrophin assembly at the sarcolemma [31]. In this context, our results validate these studies and highlight an important signaling role for *dys-1*, which evidently functions outside of its accepted role as a scaffolding protein in the DGC.

After the onset of paralysis in post-symptomatic samples, while the mitochondrial dysfunction is still present, we detect a second pathway, which perhaps tries to actively compensate for the loss of muscle structure by overexpressing genes involved in muscle formation. We do not know if these transcripts are indeed carrying out a compensatory mechanism, and more experiments need to be performed to further validate this finding. Of note, many of them were able to enhance paralysis in our RNAi experiments in **Fig. 4C,** suggesting that they are functionally translated.

Our RNAi experiments in **Fig. 4** most effectively screen for genes that play a role in the muscle and work cooperatively with dystrophin beginning early in development. The effects of RNAi knockdown are scored in the first hours after L1 animals hatch. In this light, the results of our semi permissive control experiments show that the absence of dystrophin is able to affect the very early development of the embryos that ultimately hatch with severe muscle defects. This underscores our sequencing results in PRE datasets that suggest dystrophin may play an early role, both in development and in regulating mitochondrial function, and is not simply a structural gene whose absence can impair developed muscle after continuous contraction throughout an organism’s lifespan.

The changes in gene expression observed in our POST dataset suggest that many of the genes acting as structural units within the sarcomere are involved in the progression of dystrophin-induced muscle damage (**Fig. 3**). Our RNAi experiments in **Fig. 4** confirmed the involvement of these genes in DMD progression and verify their role in a compensatory mechanism that may allow *C. elegans* to increase transcription of muscle-structure related genes in response to muscle damage.

*mup-2*, a gene that codes for the ortholog of muscle protein troponin, was able to increase incidence of muscle defects and paralysis in the dystrophin deficient *strain dys-1(cx18); hlh-1(cc561ts)*, without inducing the same effect on the control strain *hlh-1(cc561ts)*. In the same manner, the interference of *ttn-1*, an ortholog of the muscle protein titin, and *mlc-1*, an ortholog of myosin regulatory light chain, was able to increase incidence of muscle defects and paralysis in the dystrophin deficient strain *dys-1(cx18); hlh-1(cc561ts)*, without inducing the same effect on the control strain *hlh-1(cc561ts)* **(Fig. 4C)**.

Taken together these results suggest that *mup-2, ttn-1*, and *mlc-1* are all genetically connected to *dys-1*, and are not only overexpressed as transcripts when paralysis occurs, but are also expressed as proteins, as their dosage is necessary to increase paralysis in a *dys-1* dependent manner.

Outside this group of genes selected because of their role in muscle structure, we also studied upregulated genes essential in several signaling pathways. *icd-1* is the β subunit of the nascent-polypeptide associated complex. It was significantly upregulated in POST symptomatic data sets, it mediates proteins transport to mitochondria [37], and it is necessary and sufficient to suppress apoptosis [38]. *icd-1* knockdown induces a two-fold increase of incidence of muscle defect and paralysis in *dys-1* dependent manner **(Fig. 4B)**, suggesting that these two genes are genetically connected.

An observed limitation in using our approach for RNAi screens was the incidence of synthetic lethality. When combined with background mutations in *hlh-1*, several genes involved in development of muscle proved to be embryonic lethal when knocked down. Because this synthetic lethality achieved the same penetrance in single and double mutant strains, it prevented the scoring of differences between the two strains. It is important to note that this did not occur in the bulk of our experiments that knocked down muscle specific genes, meaning it is still feasible to use the method to screen the majority of genes in the genome, both muscle specific and ubiquitously expressed. This screening method is also now mature for high throughput, genome wide RNAi screens. The system can easily be adapted to 96 well, liquid screens. This can in turn provide a platform for rapidly identifying potential therapeutic targets that can be first validated in mammalian model systems, and eventually considered as part of a treatment plan for human DMD patients.

## CONCLUSIONS

We have definitely characterized that the nematode *C. elegans* fully mimics the majority of human DMD symptoms. The mitochondrial dysfunction previously observed in humans and vertebrate DMD muscle is also present in DMD worms, but is not induced by muscle fiber lesions as previously found in *mdx* mice, instead it precedes the observed muscle damage and paralysis in late stage disease phenotypes. In response to mitochondrial dysfunction, *C. elegans* activate a compensatory pathway to upregulate the expression of muscle structure related genes to stave off paralysis, which could explain their lack of a strong paralysis as observed in DMD patients.

## METHODS

### Preparation of the nematode strains

Body muscle specific PolyA-Pull expressing transgenic lines were obtained for previous crosses [18]. Young adult *C. elegans* worms were isolated on nematode growth media (NGM) agar plates seeded with OP50-1 and incubated at 31°C for 3.5 hours. Plates were then incubated for four days at 20°C and males were isolated from populations. Groups of five males were paired with 10 L4 hermaphrodites and incubated for 3 days at 20°C. Crosses between PAP and *dys-1(cx18)* and *dys-1(eg33)* strains were screened for GFP fluorescence using a Leica DM13000B microscope, and crosses between *dys-1(cx18)* and *dys-1(eg33)* strains and VS21/DM8005 strains were screened for mCherry fluorescence using a Leica MZ10F microscope. Strains positive for fluorescent markers were subjected to Sanger sequencing for the verification of mutations in the dystrophin gene using the following primers: *dys-1*(cx18)_F: GGCTTAATATGAGCTGGACGAAG, *dys-1(cx18)*_R: CGCTGTCCATCTTCTTGTGG, *dys-1(eg33)*_F: GGACGGTCATGCGACCC, *dys-1(eg33)*_R: TTTGCACACGTTGCATTTGG. In order to simplify the nomenclature, we have renamed the crossed strain *dys-1(eg33)*/PAP to DP1 and the crossed strain *dys-1(cx18)*/PAP to DP2 throughout this manuscript.

### Preparation of PRE and POST symptomatic *C. elegans* PAT-Seq Strains

*C. elegans* strains were divided into pre-symptomatic (PRE) and post-symptomatic (POST) pools using mechanical filtration using pluriStrainer cell strainers (pluriSelect). Mixed stage populations were harvested from nematode growth media (NGM) agar plates seeded with OP50-1 and pelleted at 1,500 rpm. Solid pellets of approximately 2ml were sequentially pipetted through 40 µm and 20 µm nylon cell strainers to isolate an L3/L4/Adult population and an embryo/L1/L2 population, with the 40 µm strainer retaining adult enriched populations, which were combined with the L3/L4 populations retained by the 20 µm strainer.

### RNA Immunoprecipitation

*C. elegans* strains used for RNA immunoprecipitations were maintained at 20°C on nematode growth media (NGM) agar plates seeded with OP50-1. Populations were passaged until a solid 1 ml pellet for mixed stage IPs and a solid 2 ml pellet for split stage IPs was obtained after centrifugation at 1,500 rpm. Harvested worms were then suspended and crosslinked in 0.5% paraformaldehyde solution for one hour at 4°C. Worms were pelleted at 1,500 rpm, washed with M9 buffer, and flash frozen in an ethanol-dry ice bath. Pellets are thawed on ice and suspended in 2 ml of lysis buffer (150 mM NaCl, 25 mM HEPES, pH 7.5, 0.2 mM dithiothreitol (DTT), 30% glycerol, 0.0625% RNAsin, 1% Triton X-100) [18]. Lysate was then subjected to sonication for five minutes at 4°C (amplitude 20%, 10 sec pulses, 50 sec rest between pulses) using a sonicator (Fisher Scientific), and centrifuged at 21,000 x g for 15 min at 4°C. 1 ml of supernatant was added per 100 μl of Anti-FLAG^®^ M2 Magnetic Beads (Sigma-Aldrich) and incubated overnight at 4°C in a tube rotisserie rotator (Barnstead international). mRNA immunoprecipitations were carried out as previously described [18, 19]. Total precipitated RNA was extracted using Direct-zol RNA Miniprep Plus kit (R2070, Zymo Research), suspended in nuclease free water and quantified with a Nanodrop^®^ 2000c spectrophotometer (Thermo-Fisher Scientific).

### cDNA Library Preparation and Sequencing

We prepared a total of 16 cDNA libraries from *dys-1(eg33)* mixed stages, DP1 mixed stages, DP1 PRE1, DP1 PRE2, DP1 POST1, DP1 POST2, DP2 mixed stages, DP2 PRE1, DP2 PRE2, DP2 POST1, DP2 POST2, PAP mixed stages, PAP PRE1, PAP PRE2, PAP POST1 and PAP POST2. Each cDNA library was prepared using 100 ng of precipitated RNAs. cDNA library preparation was performed using the SPIA (Single Primer Isothermal Amplification) technology (IntegenX and NuGEN, San Carlos, CA) as previously described [18, 19]. cDNA was then sheared using a Covaris Covaris S220 system (Covaris, Woburn, MA), and sample-specific barcodes were sequenced using the HiSeq platform (Illumina, San Diego, CA). We obtained ∼60-90M mappable reads (1×75) each dataset.

### Bioinformatics Analysis

#### Raw Reads Mapping

The raw reads were demultiplexed using their unique tissue-specific barcodes and converted to FASTQ files. Unique datasets were then mapped to the *C. elegans* gene model WS250 using the Bowtie 2 algorithm) [39] with the following parameters: --local -D 20 -R 3 -L 11 -N 1 -p 40 --gbar 1 -mp 3. Mapped reads were further converted into a bam format and sorted using SAMtools software using generic parameters [40].

#### Cufflinks/Cuffdiff Analysis

Expression levels of individual transcripts were estimated from the bam files by using Cufflinks software [41]. We calculated the fragment per kilobase per million base (FPKM) number obtained in each experiments and performed a pairwise with other tissues using the Cuffdiff algorithm [41]. We then used the median FPKM value >=1 between each replicate as a threshold to define positive gene expression levels. The results are shown in **Additional File 1: Tables S1-S3**. **Additional File 1: Table S4** was compiled using scores produced by the Cuffdiff algorithm [41] and plot using the CummeRbund package.

### Survival Curves

Survival curves were performed as previously described [42]. Briefly, we prepared special NGM plates each containing 330 μl of 150 mM FUDR (Sigma Life Sciences, Darmstadt, Germany). These plates were seeded with OP50-1 UV inactivated prior to plating worms to minimize contamination. We plated 25 L4 worms from each strain per plate, each across 3 replicate plates, and kept these plates in 18C incubators for the duration of the experiment. For each time point, the plates were recovered, and worms were visually inspected and counted directly in the plate using a dissection stereomicroscope (Leica S6E) and a common cell counter. The strains were scored for survival every 48 hours, with survival being defined as pharyngeal pumping or the ability to move the head in response to prodding by a worm pick.

### RNAi screens at semi-permissive temperatures

RNAi screens were performed using the temperature sensitive strains PD4605 *(hlh-1(cc561))* and LS587 *(hlh-1(cc561); dys-1(cx18))*. Strains were synchronized with bleach as previously described [43] and eggs were incubated at permissive temperature (15**°**C) until populations reached young adult stage. Young adults were then plated on NGM plates seeded with OP50-1 with 5 worms per plate and incubated at semi-permissive temperature (18**°**C) for 24 hours to allow young adults to lay eggs. Plates were then recovered, and adult worms were removed, and plates containing eggs were returned to semi-permissive temperature to incubate for 24 hours. Plates were then scored for gross defects in body morphology and paralysis. We scored approximately 100 worms per plate across 5 replicate plates for each strain and each gene tested [44].

## ACKNOWLEDGMENTS

### Funding

This work was supported by the NIH grant 1R01GM118796.

### Author contributions

We thank Stephen Blazie for the establishment of the PAP strains used for crosses and sequencing controls in this manuscript. We thank Kasuen Kotagama for assistance in optimizing the PAT-Seq protocol. We thank Shannon O’Brien for assistance in RNAi experiments.

### Competing interests

The authors declare that they have no competing interests.

## DATA AND MATERIALS AVAILABILITY

Raw reads were submitted to the NCBI Sequence Read Archive (http://trace.ncbi.nlm.nih.gov/Traces/sra/). The results of our analyses are available in Excel format as **Additional File 1: Table S3**.

## DESCRIPTION OF ADDITIONAL DATA FILES

**Additional File 1: Figure 1:** Kaplan-Meier Survival Analysis. **Figure S2 :** Sequencing results. **Figure S3:** Validation and comparison with other studies. Differential gene expression analysis in PRE-and POST-symptomatic strains. **Figure S5:** Spearman correlation of read counts in DP1 (cx18) and DP2 (eg33) PRE and POST datasets. **Table S1**: Summary of results after deep sequencing. **Table S2:** Summary of sequencing results after mapping genes to WS250.

