## Supplementary Figures for "Consecutive signaling pathways are activated in progression of Duchenne muscular dystrophy in *C. elegans*"

Additional File 1 for:

**This PDF includes:**

**Figs. S1-S5**

**Table S1 - Summary of results from PAT-Seq after deep sequencing.**

**Table S2 - Summary of sequencing results after mapping genes to WS250**

**Table of Contents**

### Additional File 1

**Fig. S1:** Kaplan-Meier Survival Analysis……………..........................................…….....3

**Fig. S2 :** Sequencing results……..........................................................................……….4

**Fig. S3:** Validation and comparison with other studies…..............................5

**Fig. S4:** Differential gene expression analysis in PRE- and POST-symptomatic strains.…….....................................................................….6

**Fig. S5:** Spearman correlation of read counts in DP1 (cx18) and DP2 (eg33) PRE and POST datasets……………………………..7

**Table S1**: Summary of results after deep sequencing………………………………….....8

**Table S2:** Summary of sequencing results after mapping genes to WS250…...........................................................................................9

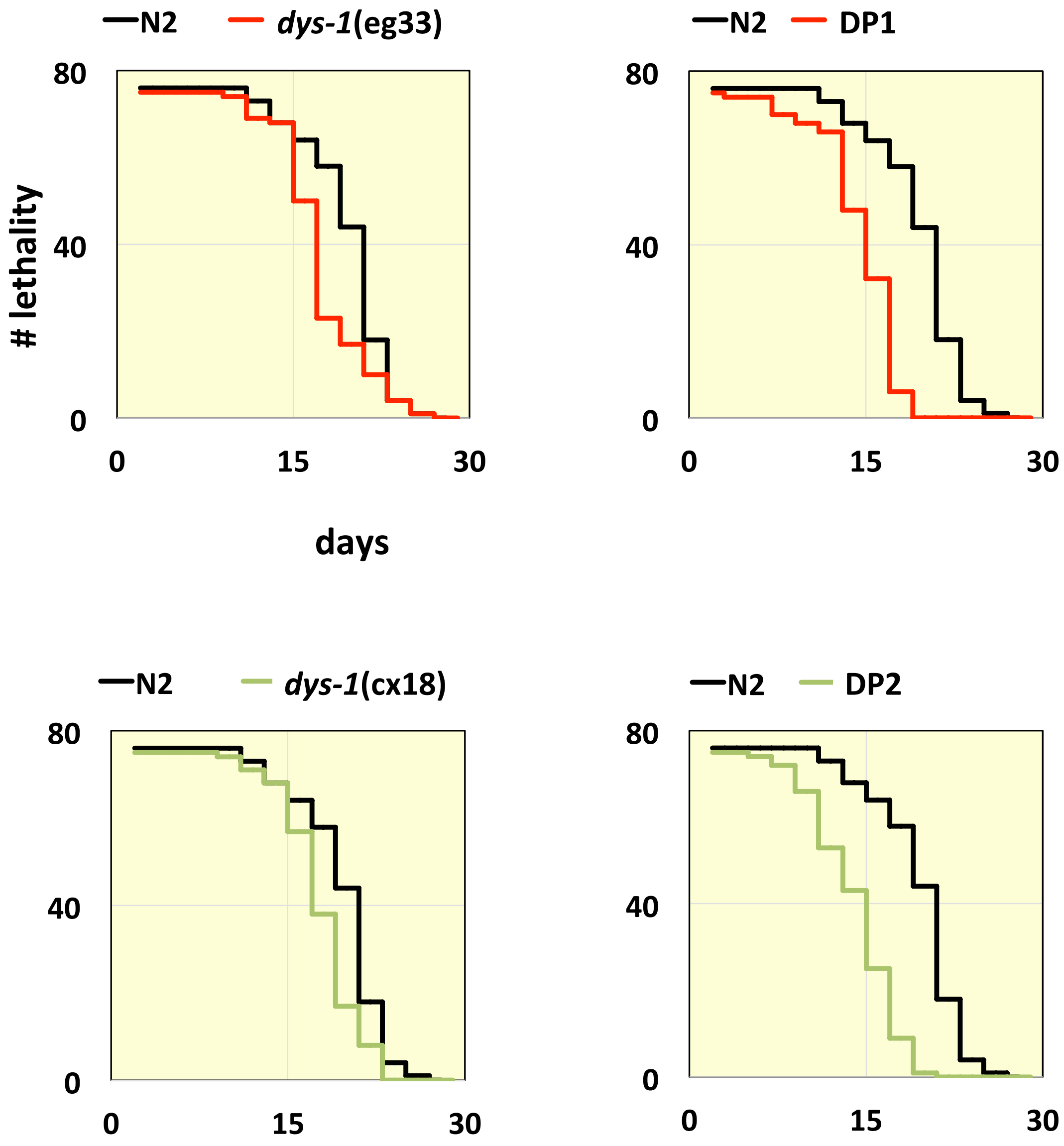


**Fig. S1**. Kaplan-Meier Survival Analysis. We have performed survival curves on *dys-1*(cx18), *dys-1*(eg33), DP1 and DP2 in order to confirm the effects of dystrophin deficiencies and the PAP cassette on average and maximum lifespan. Strains were bleached and synchronized, worms were grown to early adulthood and 25 worms per plate across three replicates were plated on NGM plates containing 150 mM FUDR. Plates were scored for survival every 48 hours. We define survival as a complete lack of movement and pharyngeal pumping after disturbance with a hair pick. Each graph represents the average percent lethality for 75 worms per control and experimental strain. Left Panels compare both dystrophin-deficient strains to wild type strains. Right Panels compare both dystrophin deficient strains after crossing with *wt* strains containing our PolyA-pull (PAP) constructs.

Fig. S2

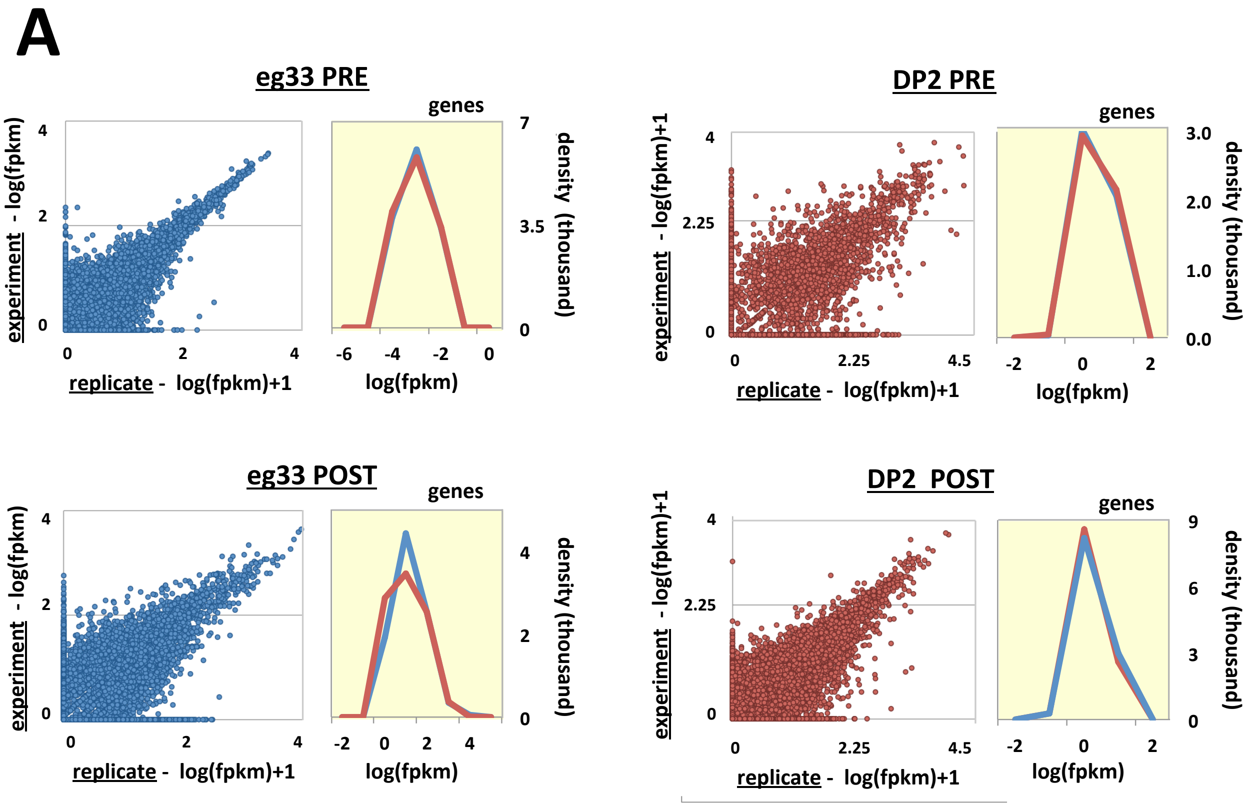


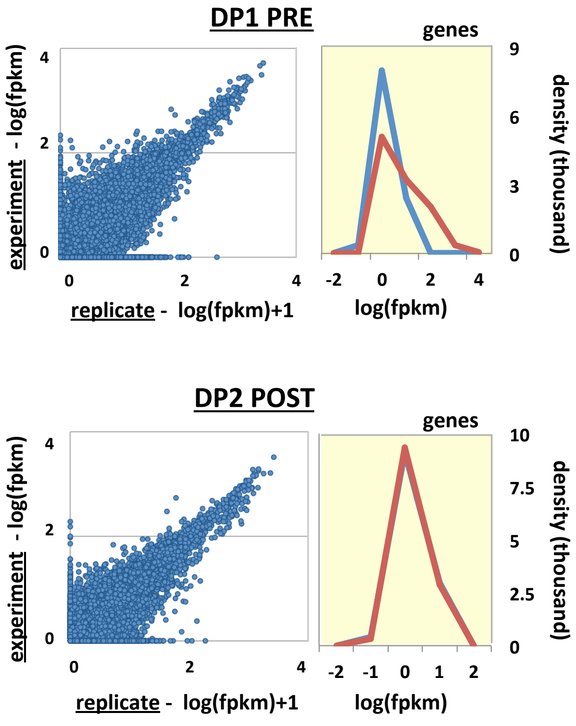


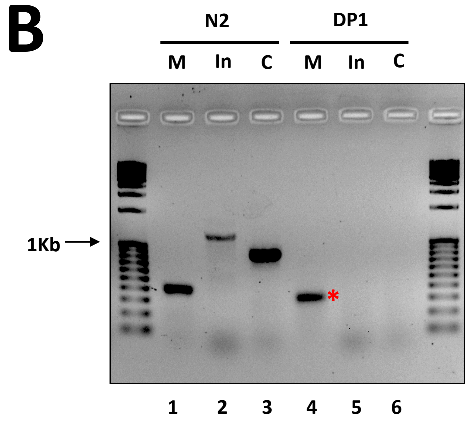


**Fig. S2: Sequencing results. (A)** Left panels: Scatter plot of mapped genes from eg33 PRE and POST, DP1 PRE and POST and DP2 PRE and POST datasets displayed by fpkm value detected in each replicate on a logarithmic (log10) scale, to highlight similarity of detection between replicates. Right panels: The distribution of the fpkm values in experiment (red) and replicate (blue) samples for each dataset. The plots were generated using the cummeRbund package v. 2.0. (B) Quantification of the specificity and sensitivity of the pull-down experiments using genomic PCR (lanes 1, 2 and 3) and RT-PCR (lane 4, 5 and 6). Using immunorecipitation we successfully isolate the muscle specific gene *myo-3* (lane 4) (*) from RNA from our DP1 strain, but not the intestine specific gene *ges-1* (lane 5) and the cuticle specific gene dpy-7 (lane 6). Lane 1, 2 and 3 shows the expected band sizes of the genes tested.

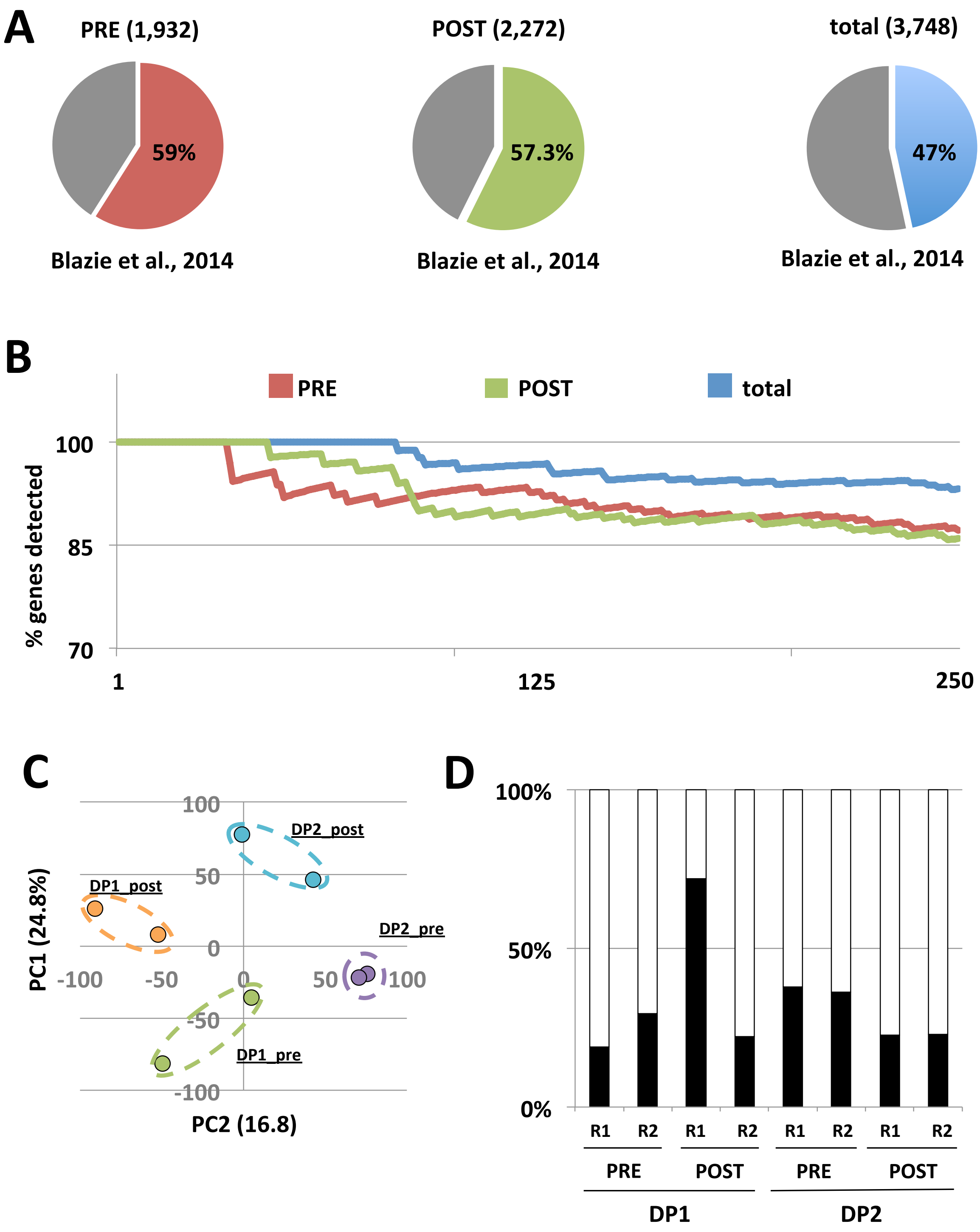


**Fig. S3:** Validation and comparison with other studies (A) Comparative analysis of genes detected in our study versus body muscle specific datasets from Blazie *et al*., 2014. The number of genes overlapping our PRE, POST and a combined dataset (total) are summarized. (B) Genes identified in this study and the muscle specific dataset from Blazie *et al*., are ranked by fpkm value and the top 250 genes from both datasets are compared. (C) Principal Component Analysis (PCA) shows high correlation among each duplicate within our datasets. (D) For this study we have used the top ~30-40% positive hits produced by Cufflinks. Black bars: % of reads used for each dataset in this study. White bars: % of reads discarded. R1 and R2: Replicate 1 and Replicate 2.

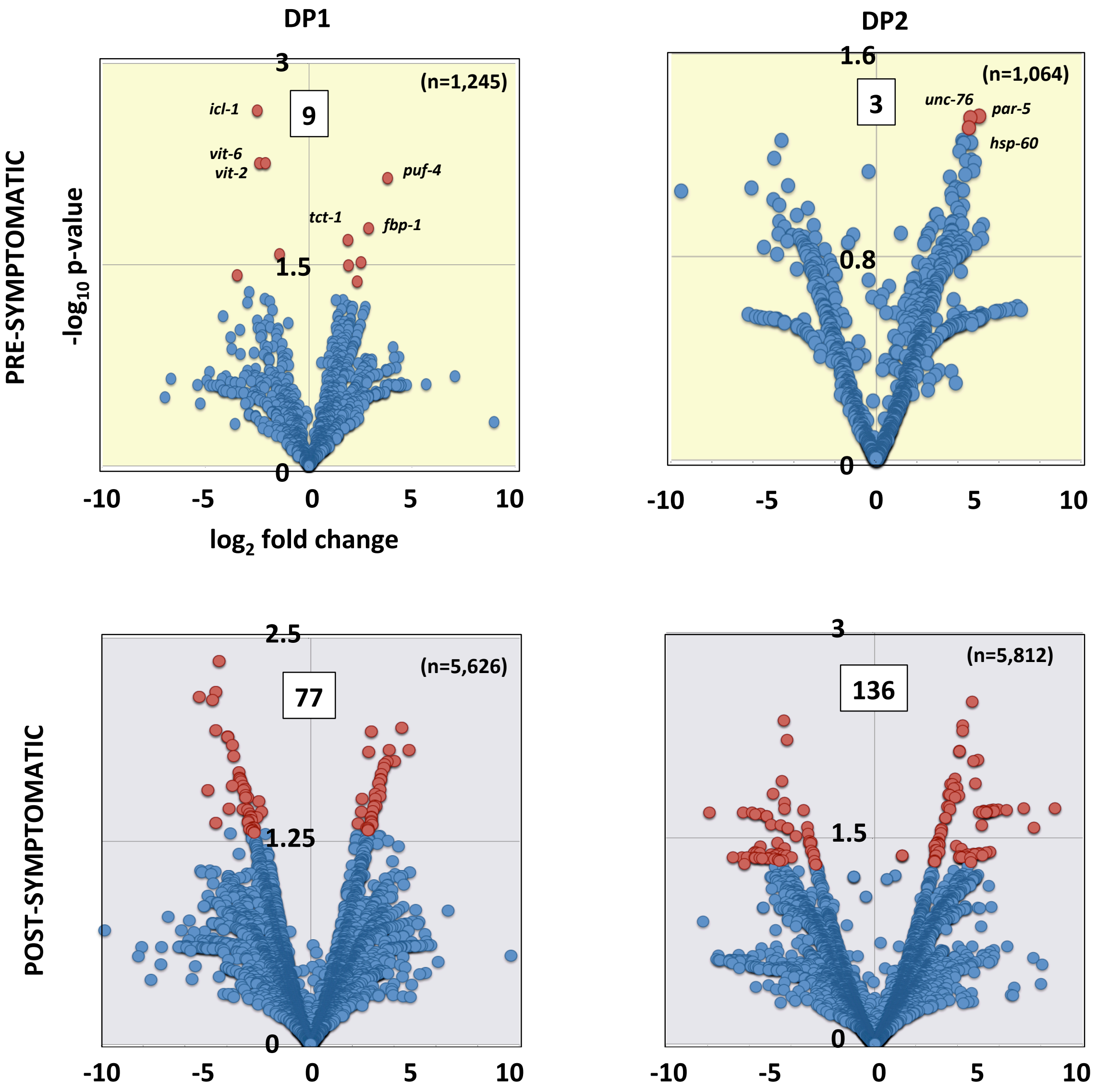


**Fig. S4:** Differential gene expression analysis in PRE- and POST- symptomatic strains. We have studied the changes in gene expression for genes detected in both PAP, DP1 and DP2 strains. The volcano plots show the changes of gene expression between each tissue (p-value versus fold-change). Total number of genes that significantly switch between PRE and POST symptomatic ($p<0.05$) are shown in red and boxed. The top genes identified in our PRE dataset are named in the chart. The density for the genes detected in the POST dataset prevents individual labeling. The complete list of genes is shown in Supplementary Table S4.

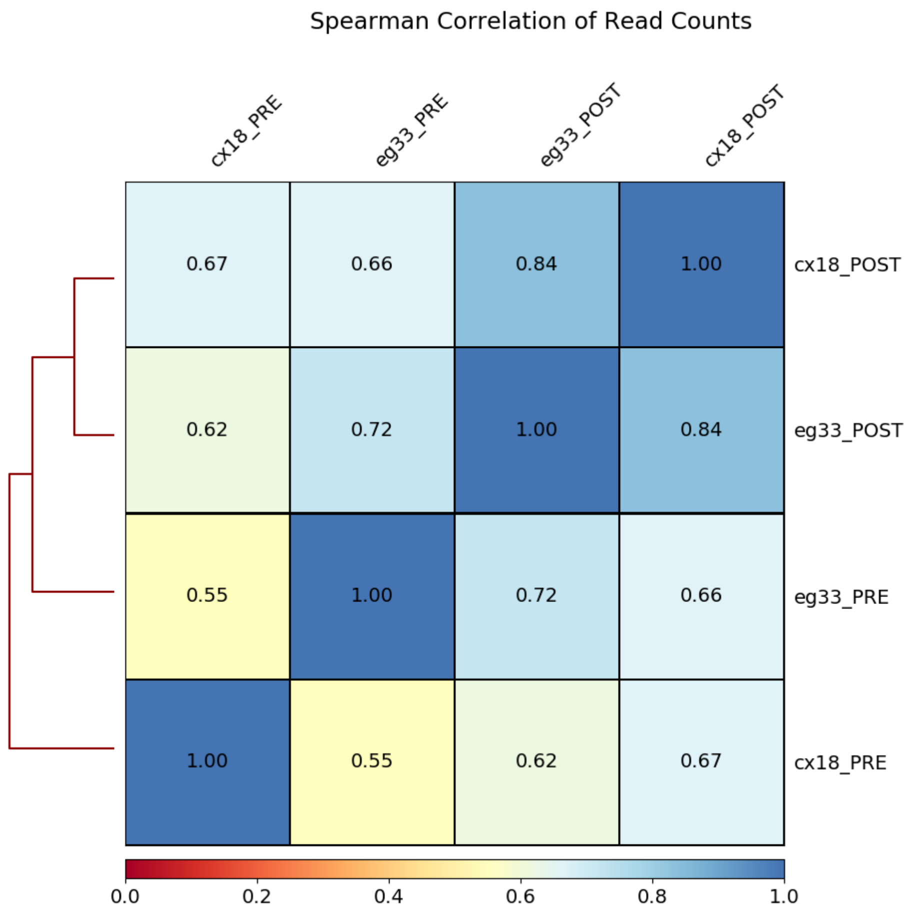


**Fig. S5:** Spearman correlation of read counts in DP1 (cx18) and DP2 (eg33) PRE and POST datasets. The chart shows an overall similarity between the DP1 and DP2 datasets. This analysis was performed using the DeepTools 2.1.0 suite (Ramirez et al., 2014).

| Sample | | | Total reads | Mapped (%) | Not mapped |
|---|---|---|---|---|---|
| *dys-1(eg33)* | PRE | experiment | 61,331,245 | 16,458,692 (27) | 44,872,553 |
| | | replicate | 46,677,723 | 17,812,390 (38.1) | 28,865,333 |
| | POST | experiment | 60,617,365 | 55,397,232 (91.39) | 5,220,133 |
| | | replicate | 45,052,554 | 40,638,981 (90.2) | 4,413,573 |
| PAP | PRE | experiment | 60,136,874 | 25,824,737 (43) | 34,312,137 |
| | | replicate | 38,934,890 | 15,628,961 (40) | 23,305,929 |
| | POST | experiment | 47,138,436 | 13,403,525 (28.5) | 33,734,911 |
| | | replicate | 35,399,372 | 15,437,939 (44) | 19,961,433 |
| DP1 | PRE | experiment | 74,708,361 | 6,519,607 (9) | 68,188,754 |
| | | replicate | 80,851,313 | 8,513,602 (10.5) | 72,337,711 |
| | POST | experiment | 85,543,247 | 60,585,159 (70.8) | 24,958,088 |
| | | replicate | 76,252,221 | 49,950,579 (65.5) | 26,301,642 |
| DP2 | PRE | experiment | 86,493,693 | 16,895,535 (19.5) | 69,598,158 |
| | | replicate | 27,624,484 | 2,174,470 (8) | 25,450,014 |
| | POST | experiment | 36,061,335 | 23,851,357 (66.1) | 12,209,978 |
| | | replicate | 29,015,306 | 17,706,345 (61) | 11,308,961 |

**Table S1**: Summary of results from PAT-Seq after deep sequencing.

| Sample | | | | Genes |
|---|---|---|---|---|
| *dys-1*(eg33) | PRE | experiment | 3,394 | 2,603* |
| | | replicate | 3,416 | |
| | POST | experiment | 2,951 | 1,738* |
| | | replicate | 2,957 | |
| PAP | PRE | experiment | 2,864 | 1,712* |
| | | replicate | 2,996 | |
| | POST | experiment | 3,487 | 2,187* |
| | | replicate | 3,353 | |
| DP1 | PRE | experiment | 2,078 | 1,100* |
| | | replicate | 2,176 | |
| | POST | experiment | 3,020 | 1,950* |
| | | replicate | 2,636 | |
| DP2 | PRE | experiment | 2,415 | 1,640* |
| | | replicate | 2,454 | |
| | POST | experiment | 3,005 | 2,266* |
| | | replicate | 2,885 | |

**Table S2**: Summary of genes detected in this study. Raw reads derived from the mRNA libraries on the Illumina Hi-Seq Instrument, mapped to the C. elegans WS250 genome annotation. Genes marked with an asterisk correspond to genes detected in both biological replicates (fpkm>=4)
